# Positive outcomes of COVID-19 research-related gender policy changes

**DOI:** 10.1101/2020.10.26.355206

**Authors:** Holly O. Witteman, Jenna Haverfield, Cara Tannenbaum

## Abstract

The COVID-19 pandemic has exposed and exacerbated gender biases in science, technology, engineering, mathematics, and medicine. Accumulating evidence suggests that female scientists’ productivity dropped during the initial lockdown period. With more time being spent on caregiving responsibilities, women may be struggling to collaborate on grant applications and launch new experiments. Scientists with disabilities or who belong to Indigenous nations or communities of color may have less time to devote to research due to health, family, or community needs. Collateral damage in this situation, the appropriate integration of sex, gender, and other identity characteristics in research content may also suffer. Sex and gender are better attended to when female scientists form part of the research team. Research funding agencies have a role to play in mitigating these effects by putting in place gender equity policies that support all applicants and ensure research quality. Accordingly, a national health research funder implemented gender policy changes that included extending deadlines and factoring sex and gender into COVID-19 grant requirements. Following these changes, the funder received more applications from female scientists, awarded a greater proportion of grants to female compared to male scientists, and received and funded more grant applications that considered sex and gender in the content of COVID-19 research. Whether or not these strategies will be sufficient in the long-term to prevent widening of the gender gap in science, technology, engineering, mathematics and medicine requires continued monitoring and oversight. Further work is urgently required to mitigate inequities associated with identity characteristics beyond gender.

## Introduction

Funding organizations, academic institutions, and peer-reviewed journals are increasingly called upon to respond to the challenges imposed by the COVID-19 pandemic on the academic community. Many researchers have seen their non-COVID-related activities slowed down or paused as the COVID-19 pandemic disrupts lives around the world, some more than others. Much of the available literature about the pandemic’s differential effects on researchers has focused on gender. During the months in which many countries imposed measures requiring people to stay home, scientists with feminine-coded names submitted substantially fewer manuscripts than scientists with masculine-coded names and were especially less represented in rapid manuscripts about SARS-CoV-2 and COVID-19 (*1–3*). This gender gap may be partly attributable to differences in time spent on professional tasks outside research, such as teaching and service, whose loads fall more heavily on white women and people of color (*3*, *4*). It may also be partly attributable to greater time spent caring for children, older adults, or disabled loved ones, responsibilities that are disproportionately borne by women (*5*). Less literature is available on identity dimensions other than gender, but it is possible that researchers belonging to minoritized groups who are over-represented among COVID-19 cases and deaths, such as racial and ethnic minorities, Indigenous Peoples, and disabled people (*6–8*), may also be shouldering additional responsibilities to students, families, or communities. Researchers with disabilities and chronic illnesses may also have additional time constraints due to the unavailability of caregivers or additional time required to minimize risks to their health (*9*).

In response to the pandemic, funding agencies around the world have rolled out rapid competitions supporting research relevant to SARS-CoV-2 and COVID-19. Because female researchers and those belonging to other minoritized groups may face additional constraints, they may be less able to quickly pivot and contribute to research in this area, even though their expertise could be needed.

## Imbalances make for less impactful science

In addition to concerns about fairness, inequities matter because funding researchers who belong to different groups helps ensure that research serves all groups. Any researcher could potentially conduct research that serves any population—and many do. Yet, for example, women are more likely to conduct health research that appropriately accounts for sex and gender (*10, 11*), and Black scientists are more likely to conduct health research that serves the needs of Black people (*12*). Failing to equitably fund researchers across groups may therefore mean short-changing the communities and groups to which they belong and which they may be well-equipped to study in nuanced, expert ways.

## Might funding agency policy changes help ensure more impactful science?

The Canadian Institutes of Health Research (CIHR), Canada’s national health research funding organization, is committed to creating an equitable, diverse, and inclusive health research enterprise. As part of this commitment, leadership regularly identifies, assesses, and implements actions to remove barriers faced by under-represented groups to access funding. In early 2020, the federal government allocated additional funding to rapidly provide resources to scientists for COVID-19 research. CIHR accordingly launched a rapid response COVID-19 funding competition in February 2020. Data-driven sex- and gender-based analyses of this first competition revealed substantially lower application pressure and success rates from the female applicant pool compared to historical funding patterns, as well as a smaller proportion of grants accounting for sex and gender in the research content. To address these problems in a second COVID-19 funding competition in April to May 2020, the organization sought input regarding potential solutions to challenges faced by female scientists with different lived experiences of career stage, caregiving responsibilities, and ability. The organization then implemented a series of data-driven gender policy interventions specific to applicants, peer reviewers, and research content (Table 1).

**Table 1:**
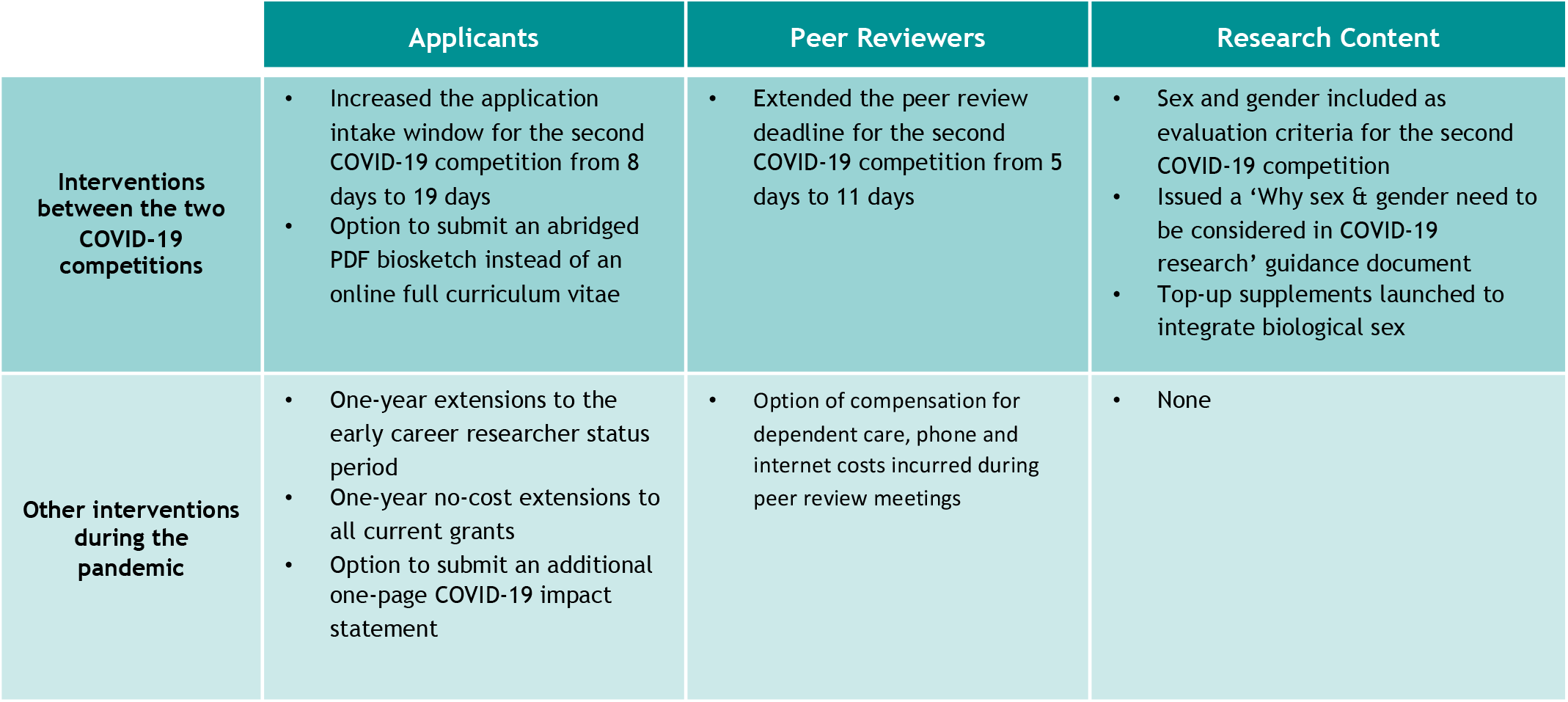
Summary of the gender policy interventions implemented by CIHR during the COVID-19 pandemic

|  | Applicants | Peer Reviewers | Research Content |
| --- | --- | --- | --- |
| **Interventions between the two COVID-19 competitions** | Increased the application intake window for the second COVID-19 competition from 8 days to 19 days<br>Option to submit an abridged PDF biosketch instead of an online full curriculum vitae | Extended the peer review deadline for the second COVID-19 competition from 5 days to 11 days | Sex and gender included as evaluation criteria for the second COVID-19 competition<br>Issued a 'Why sex & gender need to be considered in COVID-19 research' guidance document<br>Top-up supplements launched to integrate biological sex |
| **Other interventions during the pandemic** | One-year extensions to the early career researcher status period<br>One-year no-cost extensions to all current grants<br>Option to submit an additional one-page COVID-19 impact statement | Option of compensation for dependent care, phone and internet costs incurred during peer review meetings | None |

Specifically, in the second competition, CIHR provided applicants and reviewers more time and eased application requirements around investigators’ curriculum vitae. CIHR also created a guidance document titled, ‘Why sex and gender need to be considered in COVID-19 research,’ for applicants and peer reviewers, and included it as a resource to raise awareness about the importance of considering these elements in the design, methods and analysis of COVID-19 research proposals (*13*). Attention to sex and gender was made explicit in the evaluation criteria and an email was sent to peer reviewers instructing them to assess the quality and appropriateness of each proposal’s sex- and gender-based analysis. Reviewers were asked to consider and examine sex, gender, and other identity factors (e.g., age, race, ethnicity, culture, religion, geography, education, disability, income and sexual orientation) at all stages of the research process including planning and implementation of the research project and related activities.

The two COVID-19 rapid response grant competitions were otherwise similar. Both focused entirely on COVID-19, allowed research across a diverse portfolio of health research from biomedical research to population health, and allowed proposals to be five pages long with no appendices.

## Comparisons before and after policy changes

To assess the potential effects of the differences in program design between the consecutive competitions, we compared two outcomes. First, we examined application and success rates for grants submitted by principal investigators (PIs) with different identity characteristics, as well as the proportion of female peer reviewers. Second, we examined whether PIs indicated that their grants accounted for sex and gender in the web-based application form. We used grants management data routinely collected when applicants and peer reviewers create accounts in the online CIHR system and self-identify as female, male, or do not provide an entry in that field. These data were available for 100% of PIs and peer reviewers in this study across both competitions and therefore constituted the primary data for our analyses. In 2018, CIHR added another form that collects all applicants’ self-reported gender (Woman; Man; Gender-fluid, nonbinary, and/or Two-Spirit) and whether or not the person identifies as an Indigenous person (First Nations, Métis, or Inuit), as a member of a visible minority group, or as a person with a disability. All questions include the option to indicate, “I prefer not to answer.” These data were available for 82% of PIs in the first competition and 100% of PIs in the second competition. We compared data from the two COVID competitions to data from the most recent cycle of CIHR’s largest investigator-initiated open competition as a reference point. We conducted descriptive statistics and binary logistic regressions to determine the effect of integrating sex and gender on the receipt of a grant, with SPSS, version 26 (Chicago, IL, USA).

## Extra time is associated with improved numbers and success rates of female applicants

The first competition funded 100 of 227 applications for an overall success rate of 44%. The second competition garnered much larger application pressure, funding 139 of 1488 applications for an overall success rate of 9%. Table 2 shows the proportion of applicants who self-identified as female and who were successful in the first and second competitions. From the first to second COVID competition, the proportion of applications submitted by PIs who self-identified as female increased from 29% to 39%, and the proportion of successful applications with a female PI doubled from 22% to 45%.

**Table 2:** Female application pressure and success rates before and after gender equity changes to the roll out of two COVID-19 funding opportunities.

|  | Proportion of Total Applications Submitted |  | Proportion of Applications Funded |  |
| --- | --- | --- | --- | --- |
|  | Female PI | Male PI | Female PI | Male PI |
| Investigator-Initiated Open Competition | 36% (n=790) | 64% (n=1392) | 40% (n=154) | 60% (n=231) |
| 1<sup>st</sup> COVID-19 Competition | 29% (n=65) | 70% (n=159) | 22% (n=22) | 76% (n=76) |
| 2<sup>nd</sup> COVID-19 Competition | 39% (n=586) | 60% (n=898) | 45% (n=62) | 55% (n=77) |

Note: Percentages do not always add up to 100 as $\leq 2\%$ of applicants for each competition did not provide an entry in the female/male field.

PIs who self-identified as being part of a visible minority community constituted 30% of applicants in both COVID competitions (Table S1). This group of applicants comprised 28% and 26% of funded PIs in the first and second competitions, respectively. The numbers of applicants who identified as Indigenous, person with a disability, gender-fluid, non-binary, or Two-Spirit were too low in both competitions (either in submitted applications, funded applications, or both) to report in a disaggregated way. Canadian privacy norms are such that funding agencies do not report outcomes with counts under five (Table S1).

The proportion of peer reviewers who self-identified as female was 39% (n=46) in the first competition and 43% (n=397) in the second competition.

## Gender-policy interventions are associated with more frequent consideration of sex and gender in proposals

As shown in Figure 1, the proportion of applications that reported accounting for sex and gender in the proposal’s methodology increased notably between the two competitions. In the first competition, sex and gender considerations were not associated with the likelihood of receiving funding. In the second competition, applications from PIs indicating that sex was considered in the proposed work were more likely to be funded (Odds Ratio (OR) 3.13, 95% Confidence Interval (CI) 1.57-6.23). The effects of including gender were not statistically significant (OR 1.31, 95% CI 0.91-1.88).

**Figure 1:**
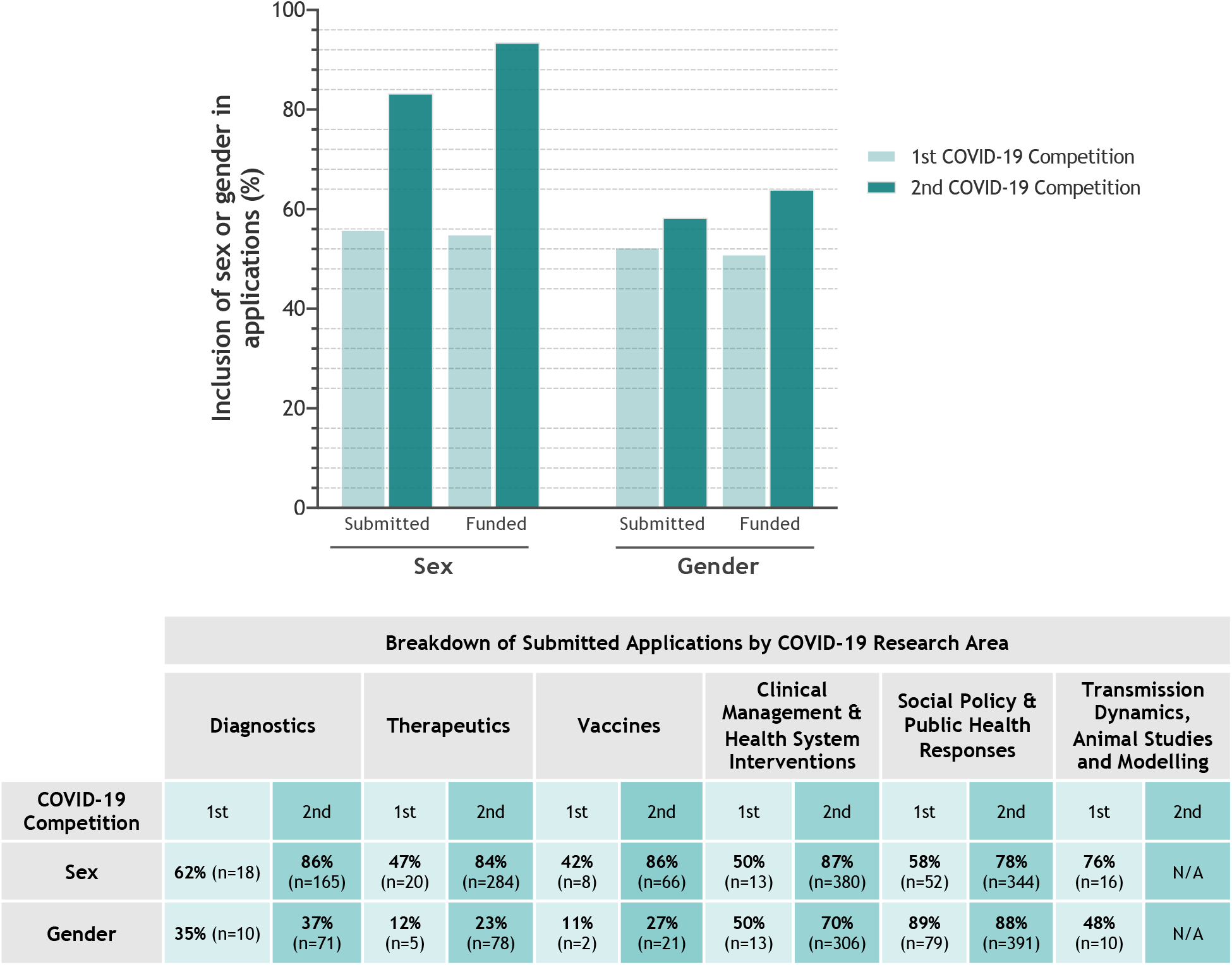
Integration of sex and gender within the research content of COVID-19 grant applications

## Opportunities for scale-up by all research funding agencies

After decades of slow progress in science, technology, engineering, mathematics, and medicine, the COVID-19 pandemic has sparked an exacerbation of existing inequities for underrepresented groups such as female scientists, racialized minorities, Indigenous individuals and individuals living with disabilities. Gender-policy interventions implemented by research funding agencies are part of a possible solution, but their effectiveness remains largely unstudied. By analyzing the outcomes of gender-policy interventions that were implemented between the first and second rounds of their COVID-19 rapid response funding competitions, CIHR offers evidence to other research funding agencies on ways to redress inequities exacerbated by the pandemic.

First, quality should not be sacrificed for speed. In the rush to launch the first COVID-19 funding opportunity, only eight days were provided to scientists to submit applications. Peer reviewers were given a similar amount of time to submit their evaluations. These short timelines may have created a disadvantage for individuals with caregiving, community, teaching, or self-care responsibilities. A relatively small increase in the application and evaluation windows was associated with a clear increase in the number of proposals submitted by female scientists. Longer intervals between the launch and submission/evaluation dates for funding competitions, as well as pre-announcements, can help provide applicants with the time required to shift commitments, plan team meetings, hold community engagement events, or set aside writing periods.

Second, implementing changes after discovering a problem is good, but it would be better to prevent problems in the first place. Because of the imbalance between male and female PIs in the first competition, the overall balance across both competitions remains out of proportion compared to historical funding patterns. This imbalance in funding may show up as differences in productivity and impact in the years to come and will need to be accounted for in evaluations of researchers.

Third, the changes implemented by the CIHR appeared to primarily solve problems related to applicants’ sex and gender. Barriers related to other identity dimensions (race and ethnicity, Indigenous identity, disability) require further analysis and consultation to identify solutions. It may more challenging to identify barriers and solutions with groups that are smaller in number, but doing so is crucial to ensure that publicly-funded research is allocated fairly and in a way that equitably serves all the public. The suite of interventions described in Table 1 provides an initial foundation to build on to advance science policy for under-represented applicants.

Fourth, educational support and explicit evaluation criteria related to sex, gender, and other identity characteristics may help ensure that applicants and peer reviewers attend to these factors within the proposed research. Such methodological rigor promotes reproducible, responsible science that benefits everyone (*14*). In the second COVID competition, both applicants and peer reviewers paid more attention to sex in the content of the research, and this was associated with a higher likelihood of funding.

It will be difficult, if not impossible, to isolate and attribute causality to each and every action put in place by funding agencies to address inequities. However, perfect cannot be the enemy of good. In the real world, during the complex circumstances surrounding the pandemic, gender policy interventions implemented by a national funding agency increased the number and success of female investigators in funding competitions and raised attention to sex and gender considerations in the science. These interventions are an important step towards dismantling systemic barriers and building better and fairer systems through and beyond the pandemic.

## Acknowledgments

We gratefully acknowledge the support of Tammy Clifford, VP Research Programs at CIHR. Adrian Mota, Sarah Viehbeck and their incredible teams helped design and implement the interventions. Christine Chambers drove implementation of the peer-review support program for evaluators with dependents.

## Author contributions

HW, JH, CT contributed to the design of the study. JH, CT contributed to data collection. JH conducted data analysis and interpretation. HW drafted the first version of the article with early revision by JH and CT. HW, JH, CT critically revised the article and approved the final version for submission for publication. JH and CT had full access to all the data in the study. HW and CT had final responsibility for the decision to submit for publication.

## Data and materials availability

Data are confidential due to Canadian privacy legislation. Researchers interested in addressing research questions related to grant funding may contact the CIHR at.

## Ethics Approval, Consent to Participate and Consent for Publication

The views expressed in this paper are those of the authors and do not necessarily reflect those of the CIHR or the Government of Canada. Data were held internally and analyzed by staff at the CIHR within their mandate as a national funding agency. Research and analytical studies at the CIHR fall under the Canadian Tri-Council Policy Statement 2: Ethical Conduct for Research Involving Humans (available: pre.ethics.gc.ca/eng/policy-politique/initiatives/tcps2-eptc2/Default/, accessed 2017 July 13.) This study had the objective of evaluating CIHR’s Investigator-Initiated programs, and thus fell under Article 2.5 of TCPS-2 and not within the scope of Research Ethics Board review in Canada. Nevertheless, applicants were informed through ResearchNet, in advance of peer review, that CIHR would be evaluating its own processes. All applicants provided their electronic consent; no applicant refused to provide consent.

## Supplementary Materials

Table S1

