## Supplemental Table 1 for "Positive outcomes of COVID-19 research-related gender policy changes"

**This PDF file includes:**

Table S1

Table S1.

|  | Feb 2020 Competition (Operating Grant: Canadian 2019 Novel Coronavirus (COVID-19) Rapid Research Funding Opportunity) | May 2020 Competition (Operating Grant: COVID-19 May 2020 Rapid Research Funding Opportunity) |
| --- | --- | --- |
| Calendar days from program launch to deadline | 8 (February 10 - 18) | 19 (April 23 - May 12) |
| Requirement in the evaluation criteria to assess sex- and gender-based analyses or their exclusion in the proposed research | No | Yes |
| Integration of sex considerations in funded grants | 55/100 = 55% | 130/139 = 94% |
| Integration of sex considerations in unfunded grants | 72/127 = 57% | 1109/1349 = 82% |
| Integration of gender considerations in funded grants | 51/100 = 51% | 89/139 = 64% |
| Integration of gender considerations in unfunded grants | 68/127 = 54% | 778/1349 = 58% |
| Overall grant success rate | 100/227 = 44% | 139/1488 = 9% |
| **Grant success rates by PI’s self-reported status as female, male, or not provided** | | |
| Female | 22/65 = 34% | 62/586 = 11% |
| Male | 76/159 = 48% | 77/898 = 9% |
| Not provided | removed due to count <5 | removed due to count <5 |
| **Grant success rates by PI’s self-reported gender*** | | |
| Women | 19/48 = 40% | 62/579 = 11% |
| Men | 66/126 = 52% | 75/864 = 9% |
| Gender-fluid, non-binary, and/or Two-Spirit | removed due to count <5 | removed due to count <5 |
| Prefer not to answer | removed due to count <5 | removed due to count <5 |
| **Grant success rates by PI’s self-reported status as a member of a visible minority*** | | |
| Visible minority | 28/68 = 41% | 36/444 = 8% |
| Not a visible minority | 54/103 = 45% | 100/975 = 10% |
| Prefer not to answer | 7/14 = 50% | removed due to count <5 |
| **Grant success rates by PI’s self-reported status as an Indigenous person (First Nations, Metis, or Inuit)*** | | |
| Indigenous | removed due to count <5 | removed due to count <5 |
| Not Indigenous | 85/176 = 48% | 136/1424 = 10% |
| Prefer not to answer | removed due to count <5 | removed due to count <5 |
| **Grant success rates by PI’s self-reported status as an person with a disability*** | | |
| Person with a disability | removed due to count <5 | removed due to count <5 |
| Not a person with a disability | 83/168 = 49% | 131/1370 = 10% |
| Prefer not to answer | removed due to count <5 | removed due to count <5 |
